# A role for δ subunit-containing GABA_A_ receptors on parvalbumin positive neurons in maintaining electrocortical signatures of sleep states

**DOI:** 10.1101/2024.03.25.586604

**Authors:** Peter M. Lambert, Sofia V. Salvatore, Xinguo Lu, Hong-Jin Shu, Ann Benz, Nicholas Rensing, Carla M. Yuede, Michael Wong, Charles F. Zorumski, Steven Mennerick

## Abstract

GABA_A_ receptors containing δ subunits have been shown to mediate tonic/slow inhibition in the CNS. These receptors are typically found extrasynaptically and are activated by relatively low levels of ambient GABA in the extracellular space. In the mouse neocortex, δ subunits are expressed on the surface of some pyramidal cells as well as on parvalbumin positive (PV+) interneurons. An important function of PV+ interneurons is the organization of coordinated network activity that can be measured by EEG; however, it remains unclear what role tonic/slow inhibitory control of PV+ neurons may play in shaping oscillatory activity. After confirming a loss of functional δ mediated tonic currents in PV cells in cortical slices from mice lacking *Gabrd* in PV+ neurons (PV δcKO), we performed EEG recordings to survey network activity across wake and sleep states. PV δcKO mice showed altered spectral content of EEG during NREM and REM sleep that was a result of increased oscillatory activity in NREM and the emergence of transient high amplitude bursts of theta frequency activity during REM. Viral reintroduction of *Gabrd* to PV+ interneurons in PV δcKO mice rescued REM EEG phenotypes, supporting an important role for δ subunit mediated inhibition of PV+ interneurons for maintaining normal REM cortical oscillations.

**Significance statement:** The impact on cortical EEG of inhibition on PV+ neurons was studied by deleting a GABA_A_ receptor subunit selectively from these neurons. We discovered unexpected changes at low frequencies during sleep that were rescued by viral reintroduction.

## Introduction

GABA_A_ receptors (GABA_A_R) represent a key inhibitory signaling system in the CNS. GABA_A_Rs are heteropentameric chloride channels, and constituent subunits influence physiological and pharmacological receptor properties. GABA_A_Rs containing the δ subunit, encoded by the *Gabrd* gene, are typically localized to extrasynaptic/perisynaptic sites of certain neuron types, where they participate in tonic and slow synaptic inhibition (Nusser et al., 1998; Cope et al., 2005). By contrast, other subunit combinations mediate fast, phasic inhibition. δ containing GABA_A_Rs have been extensively studied in principal cell types that show high δ subunit expression, including hippocampal dentate gyrus granule cells (DGC) (Wei et al., 2003; Sun et al., 2004, 2018, 2020), cerebellar granule cells (CGC) (Nusser et al., 1998; Rossi et al., 2003; Rudolph et al., 2020), and thalamocortical cells (TC) (Belelli et al., 2005; Cope et al., 2005). In these cell types, receptors containing δ subunits also contain α4 (DGC & TC) or α6 (CGC) subunits, and their activities can modulate cell excitability and transitions between firing modes (Nusser et al., 1998; Sur et al., 1999; Cope et al., 2005; Glykys et al., 2008).

In addition to excitatory principal cell types, δ subunits are expressed by inhibitory interneurons (Glykys et al., 2007, 2008). In the hippocampus, parvalbumin positive (PV+) interneurons have been shown to express receptors containing a unique partnership of α1/δ subunits that provide a source of tonic inhibition in these cells (Glykys et al., 2007). Additionally, single cell transcriptomic studies from mouse cortex and hippocampus indicate that among the interneuron cell types, PV+ cells have the strongest expression levels of *Gabrd* (Yao et al., 2021). However, because PV+interneurons primarily serve to regulate and coordinate principal cell activity (Rudy et al., 2011; Tremblay et al., 2016; Pelkey et al., 2017), it is not immediately clear how the presence and modulation of tonic/slow inhibitory currents produced by δ containing GABA_A_Rs in PV+ cells may influence coordinated network activity.

In the cortex and hippocampus, PV+ interneurons play an important role for synchronizing activity of pyramidal cells that give rise to oscillations seen in local field potential (LFP) and EEG recordings *in vivo*. Peri-somatic inhibition from PV+ interneurons is critical for the generation of gamma oscillations (30-100 Hz), and selective ablation of δ subunits from these cells increases the peak frequency but not power of *in vitro* gamma oscillations generated in the CA3 region of the hippocampus (Ferando and Mody, 2015). Beyond the generation of gamma oscillations, PV+ interneuron activity in the cortex exhibits significant phase locking to theta oscillations and increases during sleep spindles (Hartwich et al., 2009; Peyrache et al., 2011; Averkin et al., 2016; Niethard et al., 2018; Brécier et al., 2022), but the impact of slow inhibition by δ-containing receptors on these population level activities has yet to be explored.

Here we report changes to spectral content and sleep related oscillatory activity following the removal of *Gabrd* from PV+ neurons (PV δcKO). Removing a source of tonic inhibition of PV+ neurons resulted in changes to spectral content of EEG in both NREM and REM states without a change in the architecture of sleep observed in the PV δcKO mice. The oscillatory frequencies affected led us to examine sleep spindles. Total number of spindles detected across the 12-hour light cycle were not altered, but individual sleep spindle events had larger amplitudes and longer durations than the those of WT littermates. Additionally, PV δcKO mice exhibited high amplitude transient bursts of activity with a peak frequency of 7 Hz that occurred primarily during REM sleep. To distinguish whether the phenotype arose from altered network maturation following removal of *Gabrd* from PV+ neurons in early postnatal life or from acute lack of tonic/slow GABA_A_ signaling in PV+ neurons during REM states, we re-introduced *Gabrd* to PV cells in δcKO mice and found rescue of the REM phenotype. We conclude that slow inhibition of PV neurons is important for the EEG structure of REM sleep.

## Methods

### Ethical Approval

All animal procedures were performed according to NIH guidelines and approved by the Washington University Institutional Animal Care and Use Committee, protocol 22-0344. Pain and suffering were alleviated with appropriate anesthesia and analgesia during surgical procedures. Animals were reared under the care of the Washington University School of Medicine Division of Comparative Medicine. Animals had *ad libitum* access to food and water throughout. Mice were euthanized at the end of studies according to NIH guidelines for minimizing pain.

### Mice

*Gabrd* floxed mice were a generous gift from Jamie Maguire (Lee and Maguire, 2013). *Gabrd*^fl/fl^ mice were crossed with PVCre animals (Jax Strain #017320) to produce the ultimate breeding pairs consisting of Gabrd^fl/fl^XPVCre^+/-^ and Gabrd^fl/fl^XPVCre^-/-^ that generated litters containing both WT and PV δcKO mice used in this study. Animal sex and number used in this study were as follows: Main Cohorts: WT – N=8 (4M/4F), cKO – N=8 (4M/4F); Viral Rescue Cohorts: PV δcKO + *Gabrd*: N=8 (5M/3M), PV δcKO + GFP: N=7 (5M/2F), PVCre^+/-^ + *Gabrd*: N= 6 (3M/3F), PVCre^+/-^ + GFP: N=6 (3M/3F).

### Verification of genomic recombination

To confirm successful recombination of the *Gabrd* allele in PV δcKO mice genomic DNA was extracted from cerebellum chosen for enrichment of PV+ cells. PCR primers designed to produce an amplification product only from the recombined *Gabrd* locus were used to confirm PV δcKO in the brain. Only mice with confirmed *Gabrd* locus genotypes from brain tissue were included in the study.

Forward 5’ – CTCCAGTTGCCAAGCCTTTA – 3’

Reverse 5’ – CCTGGCTAATCCAGAAGGAG – 3’

### Slice electrophysiology

Mice (4 – 8 weeks old) were used for slice recordings. Coronal slices at 300 µM thickness were made from frontal cortex as described previously (Lu et al., 2023). PV interneurons were labeled by fluorescent reporter tdTomato or GFP as described. Layer 2/3 pyramidal neurons were morphologically identified through their pyramidal shape and dendritic projections toward layer 1. Borosilicate glass pipettes (World Precision Instruments, Inc.) with tip resistance of 3-7 MΩ were filled with internal solution containing the following (in mM): 130 CsCl, 10 HEPES, 5 EGTA, 2 MgATP, 0.5 NAGTP, and 4 QX-314; pH adjusted to 7.3 with CsOH; 290 mOsm). To record tonic currents, cells were voltage clamped at -70 mV and 1 µM THIP was bath applied. Tonic currents were analyzed as described previously (Lu et al., 2023).

### EEG surgery

Mice were anesthetized with isoflurane (5% for induction, 1.5-2% for surgery), and mounted in a stereotactic frame (Kopf, Tujunga, CA). Bilateral holes were drilled in the skull for insertion of epidural screw electrodes for frontal (+0.7 AP, ± 0.5 ML bregma), and parietal (-2.0 AP, ± 1.5 ML bregma) electrodes. An additional screw over the cerebellum (-1.0 AP lambda) served as a common ground reference. To facilitate vigilance scoring of EEG, a single stainless steel wire was implanted in the nuchal muscle for EMG measurement. Animals were allowed to recover in their home cages for a minimum of three days before initiating EEG recordings.

### EEG recording

For the duration of the experiment mice were maintained on reverse lighting cycle. Recording sessions were initiated 2 hours prior to lights on to prevent disturbing natural sleep behaviors during 12-hour light cycle used for analysis. EEG was acquired with an OpenEphys system as previously described (Lambert et al., 2023). Up to four mice were recorded simultaneously with genotypes and sexes being distributed equally across recording chambers between recording sessions. Signals were digitized at 1000 Hz and filtered from 1 – 250 Hz with a 2nd-order Butterworth digital filter. Raw data were imported into MATLAB for further analysis.

### Sleep Scoring

EEG/EMG was scored for sleep/wake stages with the AccuSleep toolbox for Matlab (Barger et al., 2019). A subset of 15 second epochs were scored as wake, NREM, or REM totaling a minimum of 5 minutes of each state. These were manually scored and used to calibrate a pretrained network, which subsequently scored the remainder of the epochs. A minimum of three successive epochs was required to classify a state change. Scored recordings were further reviewed by an expert scorer to validate proper scoring of state transitions.

### EEG analysis

To generate average power spectra of behavioral states, EEG scored for each state was concatenated and spectra produced from 0-100 Hz with the Matlab pspectrum function. For active wake spectra only EEG segments identified by combined evidence of animal movement, EMG, and the presence of theta rhythm in the parietal electrodes were considered. Spectra were consolidated into 256 frequency bins from 0-100 Hz prior to statistical analysis.

Sleep EEG was further analyzed with the Better OSCillation (BOSC) method (Whitten et al., 2011) applied to time frequency spectrograms from a wavelet transform of the raw EEG signal combined from either 60 minutes of NREM or REM combined from the entire recording session. Continuous wavelet transforms were produced utilizing a set of 100 complex Morlet wavelets centered from 1 – 100 Hz in 1 Hz steps with wavelet cycles increasing logarithmically from 3 – 30. The peak of sigma frequency oscillations in NREM was calculated by determining the maximum P_episode_ value of the BOSC spectra below 20 Hz for each animal.

State transition triggered spectrograms were produced with multitaper methods in the Chronux Matlab toolbox (Mitra, 2007). A sliding window of 2 seconds with a 400 ms step size was applied and multi taper parameters [TW, K] = [3 5] were used to calculate a sliding fast Fourier transform to each identified state transition. State transitions for analysis were limited to those that met the following criteria of continuous states on either side of the transition points: wake to NREM and NREM to wake both required symmetric 3 minutes of behavior on either side of transition, NREM to REM required 3 minutes NREM followed by 90 seconds of continuous REM, and REM to NREM required one minute of REM prior to transitions and at least one minute of NREM following. Individual transition spectrograms were averaged for each mouse to produce peak power and peak frequency data throughout transitions. Average spectrograms for all mice in each condition were combined for the group transition spectrograms.

Sleep spindles were detected with an automated algorithm validated for detection of spindles from rodent EEG (Uygun et al., 2019). Briefly, events were classified as spindles if the cubed root mean squared envelope of the sigma frequency bandpass filtered frontal EEG surpassed a low amplitude threshold for durations between 500 ms and 10 seconds and during that period had a peak rising above a secondary threshold. Spindle amplitude was calculated from maximum peak to trough of the z-score normalized EEG during each event. Durations were determined by the amount of time the transformed signal was above the minimum amplitude threshold. Inter-spindle-intervals were defined as the time between each spindle start point.

To generate event spectra of high amplitude transient burst activity observed during REM sleep, EEG segments containing events were removed from the input to generate a power spectrum with event-free background REM. This background spectrum was used for normalization of the spectrum generated from the total REM EEG, leaving only the REM event-related spectral profile in the final spectrum.

### Viral production and delivery

The expression plasmid pAAV-CBA-FLEX-GABRD-IRES-GFP was produced by substitution of synthesized DNA containing the coding sequence of *Gabrd,* an internal ribosome entry site, and GFP into pAAV-CBA-FLEX-GFP (Addgene # 28304). PhP.eB viral particles containing expression plasmids were produced by the Washington University Hope Center Viral Vectors Core. Viral doses from 5.8 X 10^10^-1.0 X 10^11^ vg/mouse were delivered systemically with retro-orbital injections prior to EEG surgeries. Injection volumes were determined from viral titers and animals received either one or two injections over 72 hr in alternating eyes to keep each injection volume below 150 μL. Fluorescent labeling of PV+ interneurons for slice electrophysiology in PV δcKO mice was accomplished by viral injection of PHP.eB-AAV-FLEX-GFP (Addgene #28304-PhP.eB) retro-orbitally at least 4 weeks prior to preparing acute cortical slices.

### Immunohistochemistry

Mice were perfused with phosphate buffered saline prior to dissection. Fresh cerebellar tissue was harvested to isolate genomic DNA for knockout validation. After removal of the cerebellum, brain hemispheres were separated with a midline cut and one hemisphere was immersion fixed in 4% paraformaldehyde overnight at 4°C and cryoprotected for 48h in 30% sucrose at 4⁰C. Brains were frozen on dry ice, and sagittal sections of 45 μm were cut on a freezing microtome. Free-floating sections were blocked in 3% normal donkey serum, 0.3% Triton-X detergent, and 1% BSA in PBS for 1 h at room temperature. Sections were incubated in primary goat anti-GFP antibody (Abcam, ab6673) and primary rabbit anti-PV (Swant, PV28) diluted 1:1000 in block solution at 4⁰C, overnight with shaking. Subsequently, sections were washed 3 times with PBS and incubated in secondary Alexa Fluor 555 conjugated donkey anti-rabbit (Invitrogen, A31572) and secondary Alexa Fluor 488 conjugated donkey anti-goat (Invitrogen, A11055) diluted 1:500 in PBS for 2 h at room temperature, in the dark with shaking. Finally, sections were washed 3 times with PBS, and nuclei stained with Hoechst. Regions of cortex were assessed for co-labeling of viral GFP and PV immunostaining to assess cell type specificity of viral expression.

### Behavioral Tests

Locomotor activity was measured during the light cycle by a female experimenter unaware of experimental group. Locomotion was tested over a 48-hour period in transparent enclosures (47.6 × 25.4 × 20.6 cm) constructed of Plexiglas and surrounded by computerized photobeam instrumentation (Kinder Scientific). Total ambulations, rearing, time at rest, and distance traveled were analyzed. All equipment was cleaned with 2% chlorhexidine diacetate or 70% ethanol between animals.

Alternation behavior Y maze was measured by placing mice in the center of the maze containing three arms at 120 angles, each measuring 10.5 cm wide, 40 cm long and 20.5 cm deep. Mice explored the maze for 10 min, with complete entry defined when the hindlimbs had completely entered the arm. Alternation was defined three consecutive choices of different arms with no re-exploration. We measured the number of alternations and arm entries, from which the percentage of alternations was calculated.

Accelerating rotarod was tested by training mice on the 6 cm rod (5-40 rpm) for 30 min over 2 days. On the third day, mice were tested for 1 h.

Sensorimotor gating was tested with the acoustic startle/prepulse inhibition (PPI) assay. The responses to 120 dB auditory stimulus (40 ms broadband burst) and to a prepulse anticipating the startle pulse were measured. At stimulus initiation, 1 ms force readings were measured and averaged for the startle amplitude. Startle trials were preceded by 5 min white noise (65 dB). The first 5 and last 5 trials were startle pulse alone. 20 prepulses were presented at 4, 8, or 16 dB above background). Prepulse inhibition scores were calculated as %PPI = 100*(ASRstartle pulse alone – ASRprepulse + startle pulse)/ASRstartle pulse alone.

### Experimental Design and Statistical Analysis

Comparisons of tonic current induced by THIP were performed with unpaired t-tests. Changes observed in EEG power spectra, and BOSC spectra were assessed with a Two-Way ANOVA with factors for genotype/treatment and frequency, followed by Šídák’s multiple comparisons test to determine differing frequency ranges. In the case of the event spectra for the viral conditions, Dunnett’s multiple comparisons was used to compare each treatment condition to PV δcKO + GFP virus. EEG results from each electrode were treated independently for analysis due to the qualitative differences across electrodes observed in the recordings. Comparisons of summary statistics are presented as mean ± SEM, and simple comparisons between two groups were performed with unpaired t-tests as indicated. An alpha level of 0.05 was used throughout to denote statistical differences. Further details of design are given in figure legends.

## Results

### Validation of selective loss of δ-containing GABA_A_R activity in cortex

Although mice harboring conditional deletion of *Gabrd* in PV+ neurons have been previously characterized (Ferando and Mody, 2013, 2015; Lee and Maguire, 2013), neocortical circuits have not previously been investigated, and problems with the Cre/lox approach have been documented (Kobayashi and Hensch, 2013; Song and Palmiter, 2018; Luo et al., 2020). In addition to genomic validation (see Methods), we performed functional validation by recording two classes of neurons in prefrontal cortex that express δ subunits and that are expected to be differentially affected by the conditional *Gabrd* deletion approach: PV+ interneurons and layer 2/3 pyramidal neurons (Drasbek and Jensen, 2006). We identified WT PV+ cells by crossing PV-Cre mice with the Ai14 reporter line. WT PV+ cells challenged with the δ-preferring agonist THIP (1 µM) exhibited tonic currents that were blocked by the GABA_A_ receptor antagonist gabazine. By contrast, THIP currents were reduced in δcKO PV+ cells (Fig. 1A-C). In layer 2/3 pyramidal cells, THIP current density was smaller in WT layer 2/3 pyramidal neurons than in PV+ interneurons, resulting mainly from larger membrane capacitance of pyramidal neurons, but the THIP current density was similar between WT and δcKO mice (Fig. 1D-F). Taken together, the results suggest the importance/prominence of δ-containing receptors in PV+ interneurons and the selective deletion of *Gabrd* from the targeted cell type.

**Figure 1.**
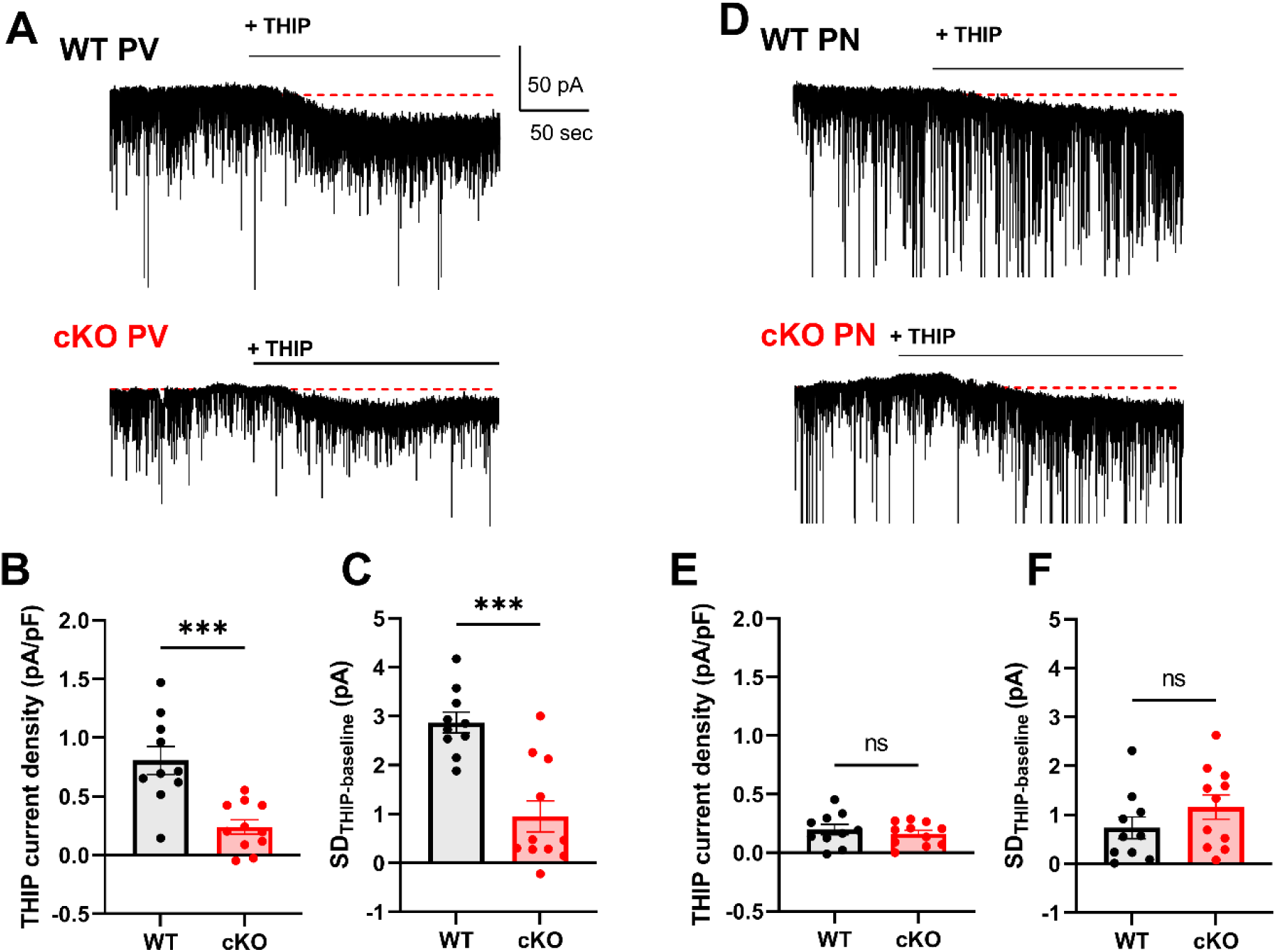
Reduced THIP currents in PV+ interneurons but not Layer 2/3 pyramidal neurons (PN) in PV δcKO Mice. **A)** Tonic currents induced by bath application of 1µM THIP recorded in PV+ interneurons in acute cortical slices from adult WT (top) and PV δcKO (bottom) animals (WT N = 10 slices/4 mice, PV δcKO N = 11 slices/4 mice). **B)** THIP current density was significantly reduced in cells from PV δcKO mice (p=0.0004, unpaired t-test). **C)** Increase in standard deviation of current induced by THIP was significantly reduced in PV+ cells from cKO slices (p=0.0001, unpaired t-test). **D)** Tonic currents induced by 1µM THIP in cortical L2/3 pyramidal neurons from WT (top) and PV δcKO (bottom) mice (WT N = 10 slices/4 mice, PV δcKO N = 11 slices/4 mice). **E)** No difference in THIP current density between L2/3 pyramidal neurons from WT or PV δcKO slices (p=0.5156, unpaired t-test). F) Increase in SD of current induced by THIP application was not significantly altered in L2/3 pyramidal neurons in PV δcKO (p = 0.2225, unpaired t-test).

### Deletion of Gabrd from PV+ cells spares sleep/wake behaviors but alters EEG spectra across sleep states

We assessed sleep/wake behaviors and network oscillations in PV δcKO mice and WT littermates with EEG across the 12-hour light phase. We scored sleep/wake behaviors of the 12-hour light cycle recording session based on parietal EEG, EMG, and movement data, categorizing states as wake, NREM sleep, and REM sleep. PV δcKO mice had similar sleep behavior patterns to WT mice (Fig. 2A), with similar distributions of wake, NREM, and REM sleep across the full recording session. Additionally, both genotypes showed a similar number of bouts of NREM and REM with comparable durations of stages (Fig. 2B-C). Taken together, the deletion of δ containing receptors from PV+ neurons did not alter the behavioral composition of sleep observed in the 12-hour light cycle.

**Figure 2.**
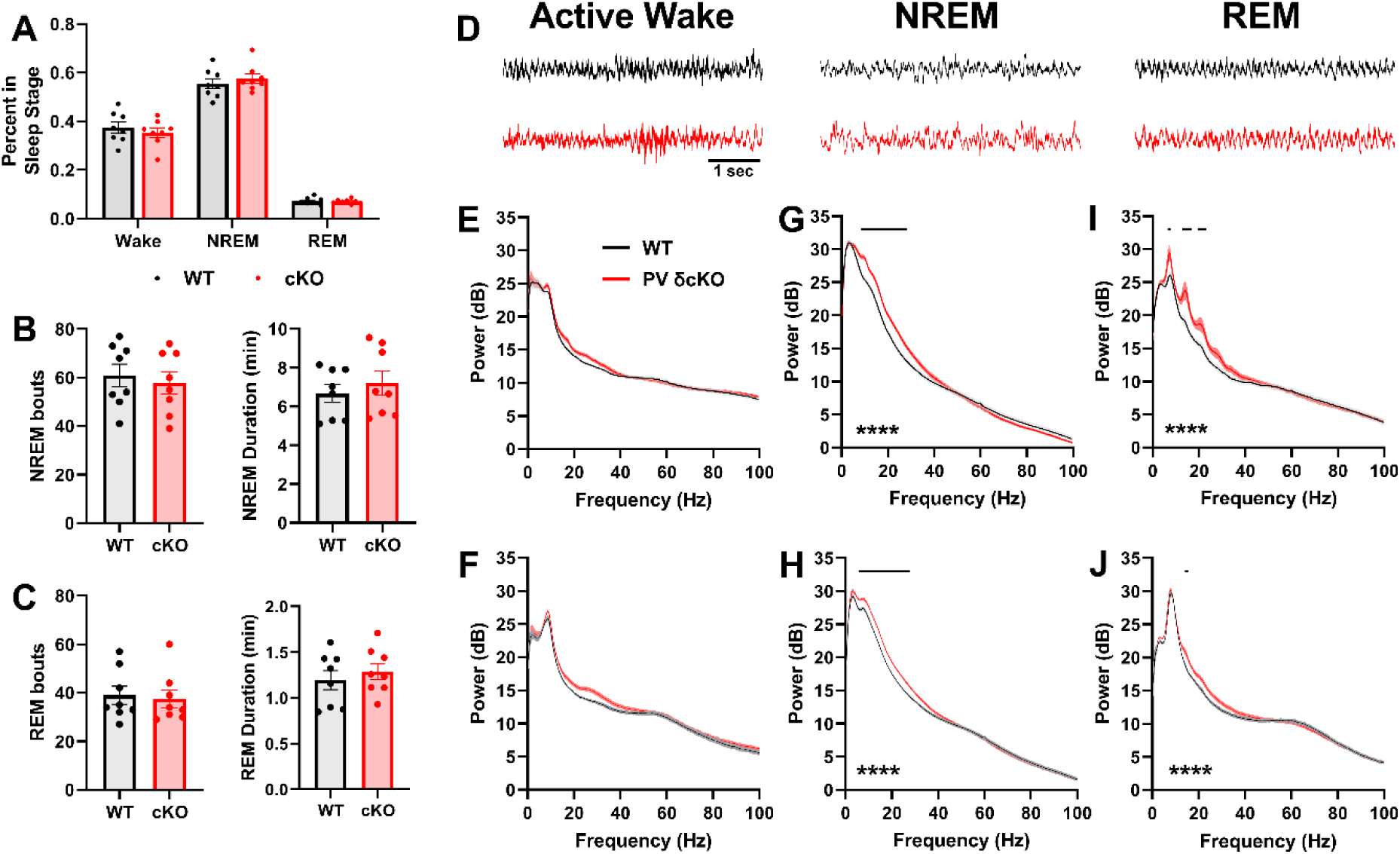
PV δcKO mice have altered EEG spectral content during sleep but no difference in sleep architecture. **A)** Proportion of 12h lights-on cycle spent in wake, NREM sleep, and REM sleep by WT and PV δcKO mice was not significantly different. **B)** Neither number of NREM bouts (left) nor average duration of NREM stage (right) was significantly altered in PV δcKO mice. **C)** Number and duration of REM bouts were similar between WT and PV δcKO mice. **D**) 5 seconds of representative Active Wake, NREM, and REM parietal EEG used for sleep scoring and initial spectral analysis from WT (black) and PV δcKO (red) mice. **E**) Frontal active wake EEG power spectra and **F)** Parietal EEG power spectra showed no significant changes in spectral content between WT and PV δcKO mice. **G)** Significant increase in the power of a broad range of low to mid frequency activity in both frontal and **H)** parietal EEG during NREM sleep (Two-Way ANOVA revealed significant Frequency X Genotype interaction in both frontal (F (255, 3584) = 5.326, p<0.0001) and parietal (F (255, 3584) = 4.268, p<0.0001) electrodes). **I)** Multi-peaked profile observed in average power spectrum of REM sleep in frontal EEG of PV δcKO mice, with significant increase in power in both frontal and **J**) parietal EEG during REM sleep (Two-Way ANOVA showed significant Frequency X Genotype interaction in both frontal (F (255, 3584) = 2.787, p<0.0001) and parietal (F (255, 3584) = 1.747, p<0.0001) electrodes). Horizontal bars in G-J indicate all frequencies with significant differences from Šídák’s multiple comparisons test (N = 8 (4M/4F) mice / genotype).

Initial EEG analysis produced average power spectra from behaviorally similar periods (Fig. 2D) across the recording session to detect any consistent changes to oscillatory activity during each behavioral state. During active wake, PV δcKO mice exhibited normal theta oscillations with no statistical difference to broad gamma frequency power in cortical EEG recordings (Fig. 2E-F). Although we observed a small increase in power in the 20-40 Hz domain, there was no Genotype X Frequency interaction detected in either frontal or parietal EEG for this behavioral state. In NREM sleep PV δcKO mice had reliably elevated power across broad frequencies spanning theta through beta ranges (Fig. 2G-H). Spectra produced from combining REM stages across the recording session showed changes more prominent in frontal EEG (Fig. 2I) but still detected in parietal EEG (Fig. 2J). Frontal EEG REM spectra in PV δcKO mice contained multiple peaks likely produced as harmonics of peak at a fundamental frequency in the theta range (Fig. 2I). The power of the first three of these frequency peaks was increased in the frontal spectra. In parietal EEG the REM spectrum was dominated by theta rhythms generated from dorsal hippocampus in both genotypes but still showed a significant Genotype X Frequency interaction across the entire spectrum. These findings suggest that signaling through δ-containing GABA_A_R in PV+ cells may regulate oscillatory activity generated in the cortex during sleep behaviors. Although we cannot exclude an effect on waking EEG (Fig. 2E, F), we focused on sleep EEG phenotypes because of the statistically robust findings and unexpected effect in lower frequency bands (Fig. 2G-J), as opposed to the gamma band, where PV+ interneurons have been most strongly implicated (Ferando and Mody, 2015; Averkin et al., 2016; Antonoudiou et al., 2020; Hadler et al., 2024).

### Changes to oscillatory activity in both NREM and REM sleep in PV δcKO mice

Although initial analysis of sleep spectra showed altered power in PV δcKO mice, these results represent the average network activity present in these behavioral states across the entire 12-hour recording session. To assess alterations to oscillatory activity during sleep stages with more temporal precision, we utilized the BOSC method that detects oscillations from a time-frequency representation of EEG and allows for both power and duration thresholds to separate oscillatory activities from background EEG. Based on observations in average spectra above, we limited BOSC analysis to frontal EEG which showed altered activity in both NREM and REM stages (Fig. 3). Additionally, focus on frontal recordings allowed us to probe transient sleep spindle activity with higher resolution during NREM (Kim et al., 2015) (see below).

**Figure 3.**
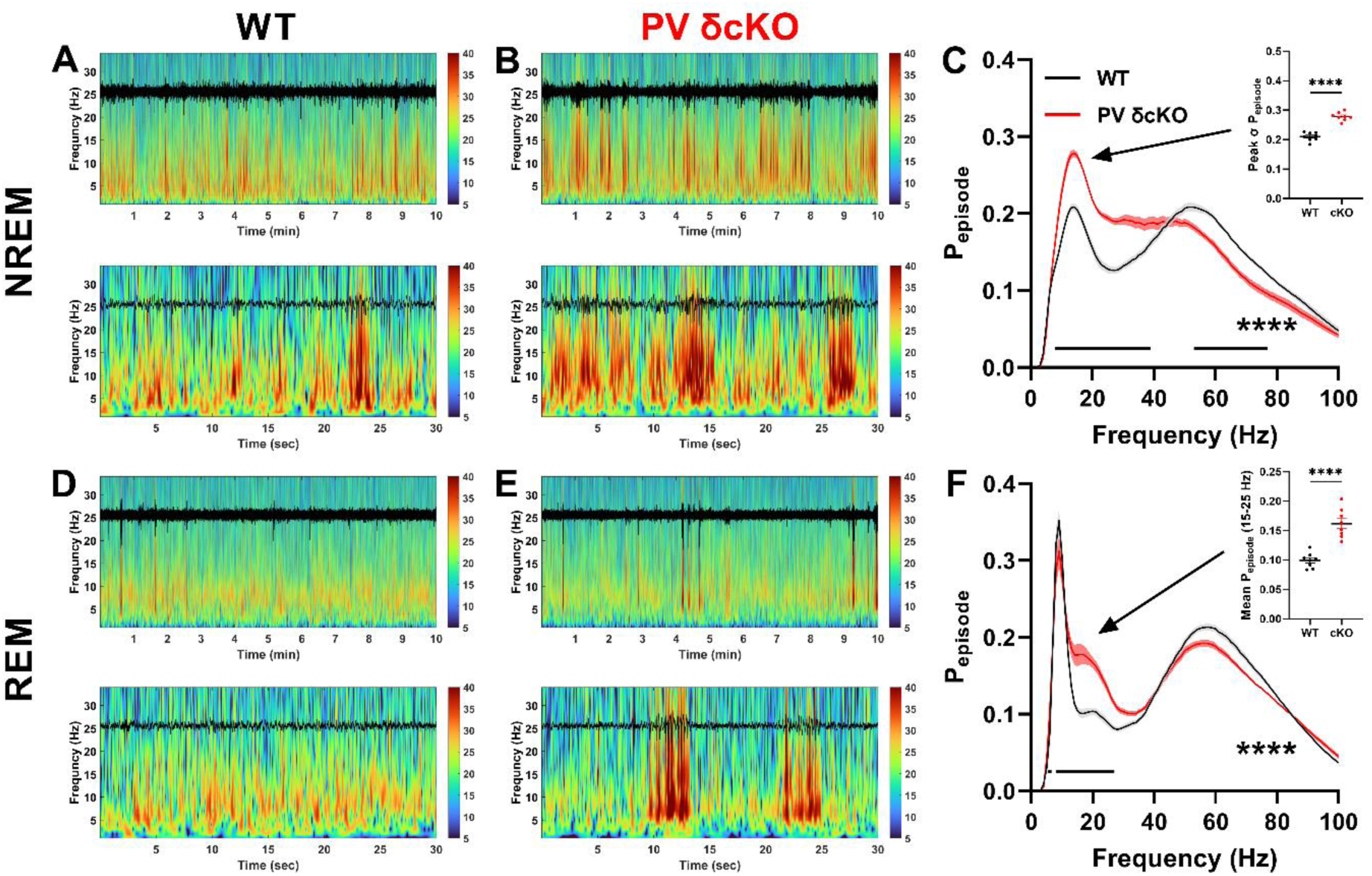
BOSC analysis of sleep stages in frontal EEG highlights increased oscillatory activities in PV δcKO mice. **A)** Continuous wavelet transform of 10 minutes NREM sleep from frontal EEG in a WT mouse (black overlaid signals throughout), lower panel shows expanded timescale of 30 seconds NREM containing oscillatory events consistent with sleep spindles. **B)** Representative wavelet transform of NREM sleep from a PV δcKO mouse showing larger periodic increases in power, expanded timescale shows longer durations of high power oscillatory activity in sigma frequency ranges. **C)** Average BOSC spectra from periods of NREM sleep show increased periodic σ frequency oscillatory activity with a peak frequency of 14 Hz *inset:* significant increase of P_epidode_ σ range peak in PV δcKO mice (p<0.0001, unpaired t-test). Two way ANOVA revealed significant Genotype X Frequency interaction (F (99, 1400) = 26.08, p<0.0001). **D)** Wavelet transform of 10 minutes combined REM sleep from a representative WT mouse showing theta frequency oscillations, expanded timescale in bottom panel. **E)** Wavelet transform of frontal EEG during REM sleep scored based on parietal signals from a representative PV δcKO mouse. Transient, high amplitude oscillatory bursts override theta frequency oscillatory activity typical of REM sleep. Expanded time scale of lower panel reveals similarity of spectral profile of transient events to periodic oscillatory activity observed in NREM sleep in both genotypes. **F)** Average BOSC spectra from REM sleep across the full recording session contains strong theta rhythm associated peak with increased oscillatory activity detected in PV δcKO mice centered around 20 Hz *inset:* significant increase in the mean P_episode_ observed from 15-25 Hz in cKO animals (p<0.0001, unpaired t-test). Two Way ANOVA indicated a significant Frequency X Genotype interaction (F (99, 1400) = 10.12, p<0.0001). Horizontal bars in panels C and F indicate all frequencies with significant differences after Šídák’s multiple comparisons test.

Continuous wavelet transforms of frontal EEG during NREM showed typical periodic increases in power spanning frequencies ranging from 5-25 Hz in both WT and PV δcKO mice (Fig. 3A-B). BOSC spectra produced from these time-frequency transforms revealed changes to NREM oscillations in PV δcKO mice with an increase in the peak P_episode_ of oscillatory activity detected in the sigma frequency range (10-15 Hz) typically associated with sleep spindle activity (Fig. 3C). Wavelet spectrograms of frontal EEG during REM states displayed concentrated power within the theta range in both genotypes (Fig. 3D-E). In PV δcKO mice, transient increases in power expanding into frequencies in the beta range were associated with instances of high amplitude bursts of activity in the raw EEG (Fig. 3E lower panel). BOSC spectra produced from these transforms showed a prominent theta frequency peak indicative of REM state with increased oscillatory activity detected in frequencies up to 27 Hz (Fig. 3F). The changes observed with BOSC based analysis confirmed that spectral alterations in PV δcKO mice are at least in part due to altered transient activities rather than to persistent changes to background EEG or stationary aspects of the EEG signals.

### NREM sleep of PV δcKO mice contains altered sleep spindles

Following observations of altered sigma frequency oscillations during NREM states in PV δcKO mice combined with previous studies showing the recruitment of cortical PV interneuron activity by ascending thalamic inputs during spindle oscillations (Hartwich et al., 2009; Niethard et al., 2018; Fernandez and Lüthi, 2020), we assessed potential alterations to sleep spindles, characterized by activity in the sigma band, in PV δcKO mice. We first examined the transition between NREM and REM sleep which is known to be enriched for sleep spindle activity (Franken et al., 1998; Astori et al., 2011; Bandarabadi et al., 2020). Average NREM to REM transition spectrograms of frontal EEG from WT and PV δcKO groups (Fig. 4A) show persistent low frequency power in NREM shifting to theta dominated REM sleep. A surge in spindle activity during the transition corresponds with a transient increase in sigma frequency power in frontal EEG in both genotypes. In PV δcKO mice, baseline NREM sigma power was elevated (Fig. 4B-C), and the peak in sigma power preceding the REM transition was higher. Despite this, the relative increase in sigma power was not different between genotypes (Fig. 4C, right). We interpret these changes to suggest that PV δcKO mice exhibit changes to spindle-like activity that is not dependent on spindle activities associated solely with state transition.

**Figure 4.**
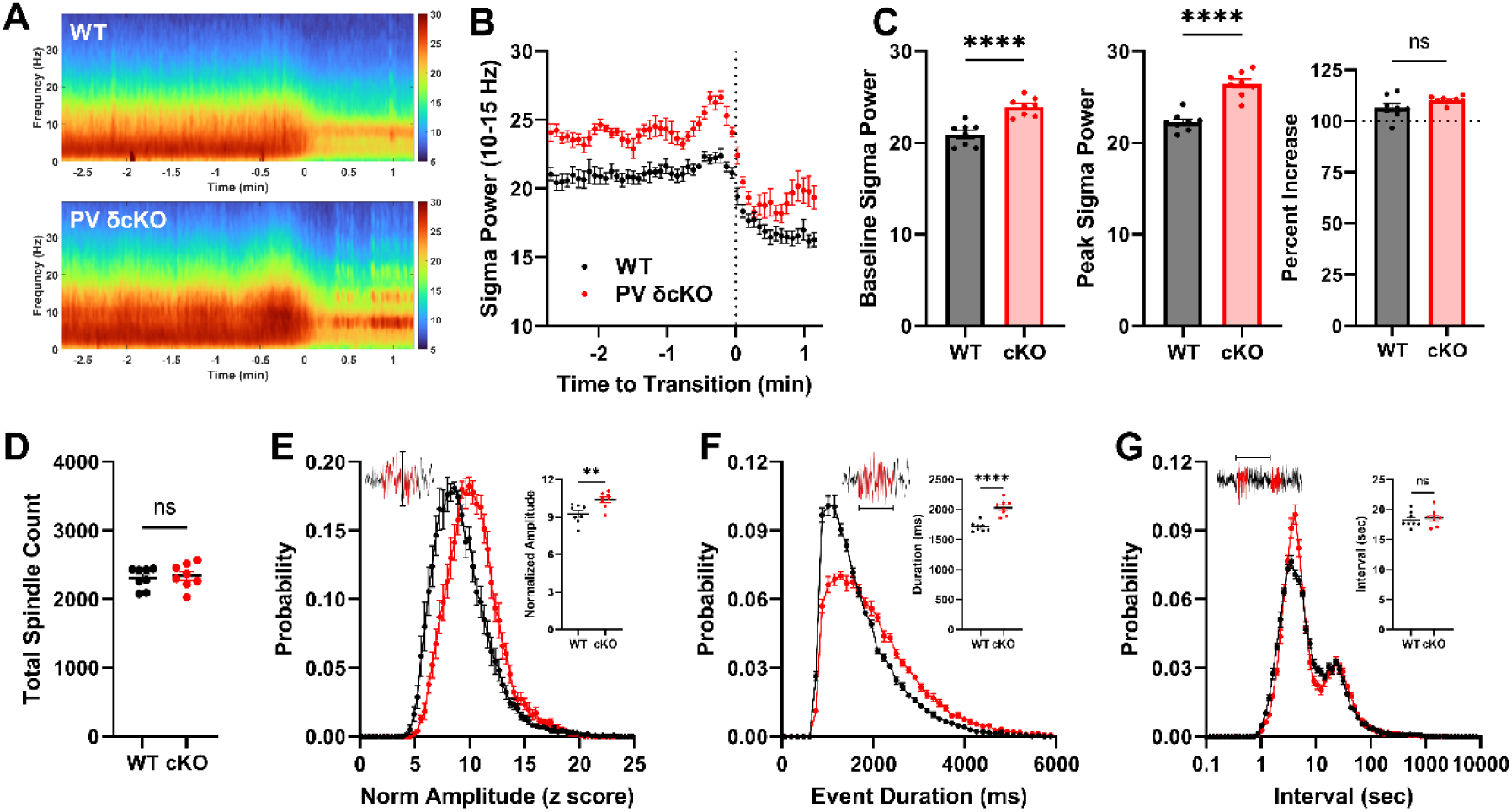
PV δcKO mice have enhanced spindle-related power at state transitions and larger sleep spindle oscillations in frontal EEG. **A)** Grand average NREM to REM state transition spectrogram from all WT (top) and PV δcKO mice showing relative increase in spindle-associated power around 10-12 Hz as the animals approach state transition, with additional evidence of altered activity after the transition to REM in PV δcKO mice. **B)** Average sigma power throughout NREM to REM transition calculated from the average transition spectrogram for each mouse. **C)** Increased baseline (before -1 min to transition) sigma power (p<0.0001, unpaired t-test), higher power sigma frequency peak during transition state (p<0.0001, unpaired t-test), but no significant difference in the relative increase above ongoing sigma frequency activity (p=0.0927, unpaired t-test) in PV δcKO mice during NREM to REM state transition. **D)** No difference in total spindle incidence from automated detection across the full recording. **E)** Spindle amplitude distributions shifted toward higher amplitudes for PV δcKO mice with inset showing significantly increased average spindle amplitude (p=0.0056, unpaired t-test). **F)** Spindle event duration distributions exhibit reduced probability of shorter duration events and increased probability density in long event duration (>2 sec) tail region, inset shows significantly increased average event duration for PV δcKO mice (p<0.0001, unpaired t-test). **G)** Start to start interval distributions show little difference between genotypes, with both groups possessing two main peaks corresponding to the periodic occurrence of temporally clustered spindle events, inset shows now significant difference to average inter-spindle-interval (p=0.6077, unpaired t-test).

To test this hypothesis, we used a previously validated algorithm for automated detection of sleep spindles from rodent EEG (Uygun et al., 2019). From the 12-hour recording session we found no difference in total spindle incidence between genotypes (Fig. 4D). Following detection of individual spindle events, we analyzed spindle amplitudes, spindle durations, and inter-spindle-intervals. We found that removal of δ containing receptors from PV+ cells resulted in higher spindle amplitudes and durations (Fig. 4E-F); however, spindle frequency measured by inter-spindle-intervals was unchanged in PV δcKO mice (Fig. 4G). Although the vast majority of events detected as spindles occur during NREM states (94.0 ± 1.2% WT vs. 92.5 ± 1.2% PV δcKO), PV δcKO mice exhibited a small increase in number of events detected in REM sleep (1.1 ± 0.1 % (27 ± 3 events) vs. 3.0 ± 0.6% (70 ± 15 events), p=0.0073, unpaired t-test). Because of the relative purity of abnormal events in REM of PV δcKO mice, we focused on high-amplitude events seen above in the REM wavelet spectra (Fig. 3E) for further characterization.

### High amplitude transient bursts of theta-frequency activity observed in PV δcKO mice enriched during REM

Throughout the recording sessions PV δcKO mice can be distinguished by the presence of periodic high amplitude bursts of activity primarily during sleep states and quiet wakefulness preceding transitions into NREM sleep. Although these events resemble some features of sleep spindles (frontal prominence, waxing and waning envelope, clustering of incidence) they are distinguished by their persistence during REM sleep when spindles typically subside to theta oscillations. Here, we present three examples of these events occurring in a PV δcKO mouse. When observed in quiet wakefulness preceding sleep (Fig. 5A left), high amplitude bursts of activity are associated with behavioral arrest, indicated by the abrupt cessation of movement for the two second duration of the event (accelerometer records). Additionally, we show repetitive occurrence of these events during a complete 90 second REM bout (Fig. 5A center) containing many high amplitude events. While repetitive bursts of activity were evident in frontal and parietal EEG, relative silence persisted in EMG and accelerometer records. An expanded timescale of one of these events demonstrates the background persisting theta oscillations of REM present in the parietal EEG and highlights the frontal prominence of event incidence and amplitude. A representative event occurring at a state transition from REM to NREM (Fig. 5A right) shows individual event durations can last many seconds and that events typically cease upon transition to NREM.

**Figure 5.**
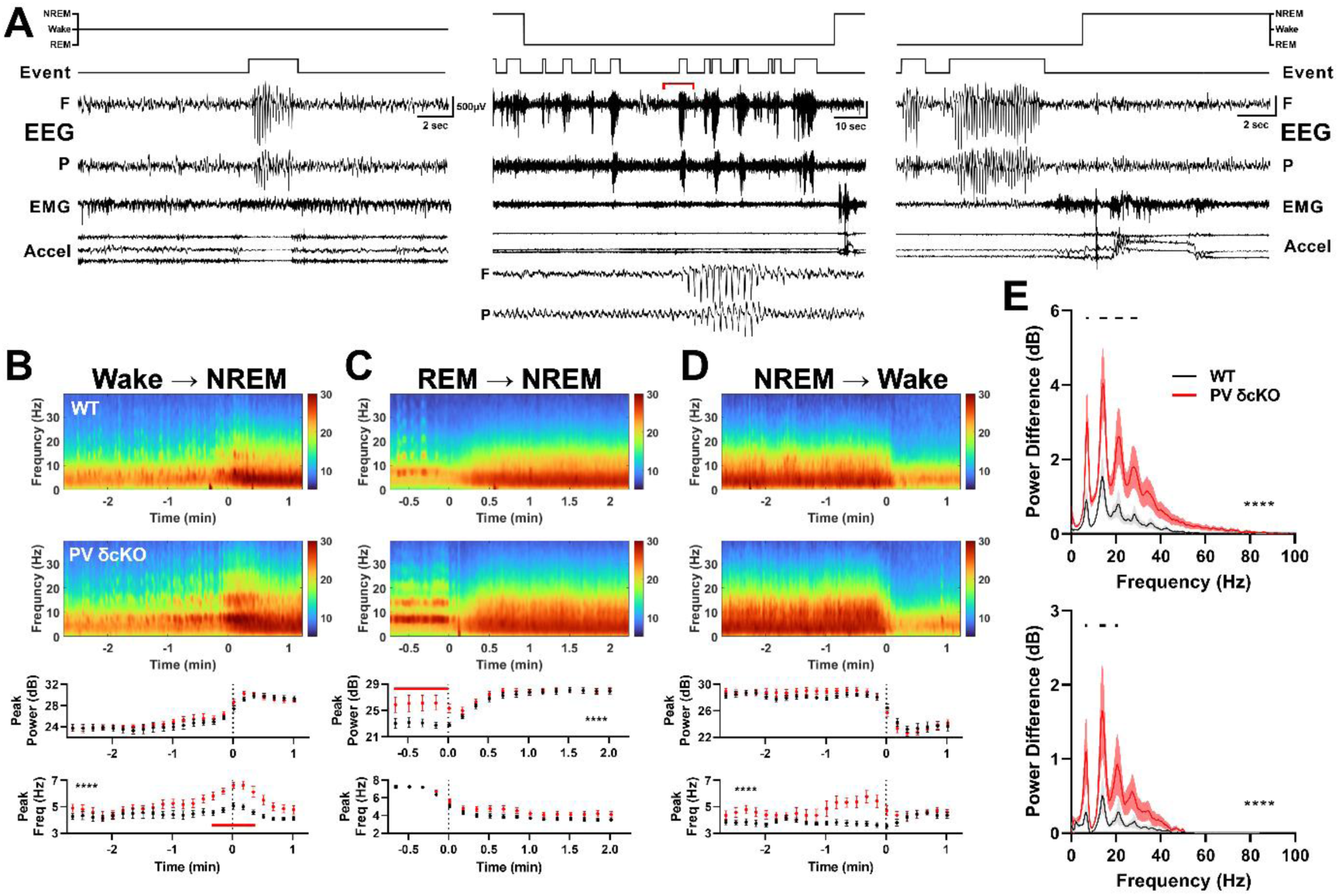
PV δcKO mice exhibit increased incidence of transient bursts of high amplitude theta-frequency activity during state transitions and REM sleep. **A)** Three representative examples of high amplitude synchronous EEG activity in a PV δcKO mouse. Records from top: hypnogram, spindle detection result, frontal (F) and parietal (P) EEG, nuchal EMG, and three-axis accelerometer. Examples include a singular event during quiet wake (left panel) in which the animal undergoes behavioral arrest for the duration of the event. The central panel displays an entire 90 second REM bout with high amplitude events increasing toward the transition from REM back to NREM. EEG for event indicated with red bracket is expanded below with strong parietal theta rhythm supporting scoring of REM brain state. The right panel presents a protracted (>4 sec) high-amplitude event immediately preceding the transition from REM to NREM sleep stage. **B)** Average spectrograms for Wake to NREM state transitions, both genotypes display a slow ramping of increased power while transitioning to NREM, with PV δcKO mice exhibiting a significantly higher frequency peak at the time of transition into NREM sleep. Two Way ANOVA reveals significant Time X Genotype interaction for the evolution of peak frequency during Wake to NREM transition (F (22, 308) = 2.875, p<0.0001). **C)** REM to NREM transitions show high incidence of high amplitude events in PV δcKO mice, with significant increase in theta frequency peak power during REM leading up to the state transition. Two Way ANOVA shows significant Time X Genotype interaction for peak power (F (16, 22F (16, 224) = 4.1934) = 4.193, p<0.0001). **D)** NREM to Wake state transitions are spared from high amplitude oscillatory bursts. Two Way ANOVA shows significant Time X Genotype interaction with respect to peak frequency during state transition to waking (F (22, 308) = 4.063, p<0.0001). **E)** Spectral profile of events occurring during REM revealed following normalization of total REM spectra to event-free REM background activity. Event frequency spectra power is more prominent in frontal (top) than parietal (bottom) electrodes. Two-way ANOVAs reveal significant Frequency X Genotype for both frontal (F (255, 3584) = 3.311, p<0.0001), and parietal (F (255, 3584) = 2.233, p<0.0001) electrodes).

To determine if events were constrained by brain state, we identified state transitions and characterized the peak frequencies observed through the transitions. In wake to NREM state transitions (Fig. 5B) we observed an emergence of some transient bursts around the transition evidenced by a transient increase in the frequency of spectral peaks in PV δcKO mice. REM to NREM transitions (Fig. 5C) were dominated by the presence of high amplitude events in REM that subsided upon transition to NREM. The presence of these events resulted in an increase in the peak power during REM preceding the transition in PV δcKO mice. NREM to wake transitions (Fig. 5D) exemplified a transition occurring in the absence of high amplitude events, but a Frequency X Genotype interaction was observed for peak frequency of the transition.

Because of the relatively high event prevalence in REM states, uniformity of REM EEG in rodents, and lack of sleep spindles contributing to periodic increases in spectral power, we focused further characterization to REM states. Removal of periods of REM EEG with events allowed the generation of a baseline REM spectrum for each animal. We then could use this baseline spectra to normalize the spectra previously generated from the entirety of REM states across the full recording session. This revealed the theoretical spectral profile of the events alone (Fig. 5E) since ongoing normal REM spectral content would be subtracted from the spectra. The spectral profile of REM events in PV δcKO showed multiple evenly spaced peaks consistent with harmonics produced from a fundamental frequency of 7Hz. The power of event spectra was greater in frontal EEG (Fig. 5E top) than parietal EEG (Fig. 5E bottom), but event related spectral power was still larger than any persisting spectra from WT mice.

### PV δcKO mice do not exhibit any behavioral abnormalities compared to their WT littermates

To further evaluate whether findings of altered EEG during sleep states could represent changes in brain activity relevant for cognitive functions we performed an initial battery of behavioral assays in PV δcKO mice and compared results with littermate controls. EEG signals of similar characteristics to those identified here (i.e., sleep spindles) have impact on motor and cognitive learning, so we focused on screens associated with these behaviors (Kam et al., 2019; Peyrache and Seibt, 2020). 48-hour spontaneous activity monitoring identified no differences in amount of ambulatory activity or rearing events between PV δcKO mice and their WT littermates (Fig. 6A, B). A potential motor coordination and/or learning deficit was followed up by testing a multi-day accelerating rotarod test (Fig. 6C) and an inverted screen test (Fig. 6D). Neither revealed a deficit in PV δcKO mice. To screen for potential deficits in working memory, spontaneous alterations were measured in a Y-maze task. The task revealed no difference in percent alterations across genotypes and no difference in the distance traveled (Fig. 6E, F). Altogether, the loss of *Gabrd* in PV neurons and subsequent changes to EEG patterns observed in sleep states did not correlate with gross changes in any behavioral parameters we assessed.

**Figure 6.**
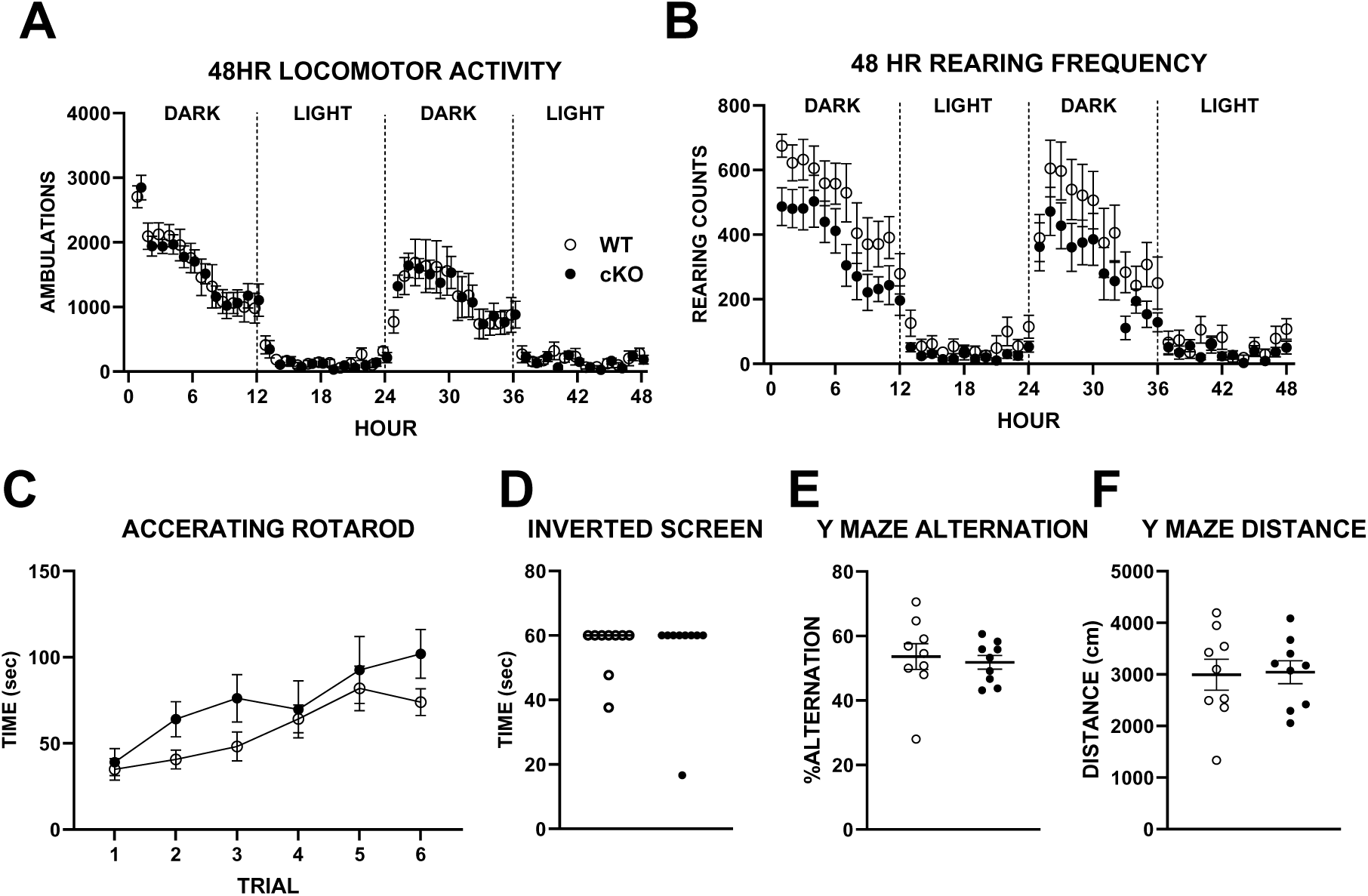
Lack of behavioral phenotype in initial battery. **A)**. Locomotor activity and **B)** rearing were measured for 48 h as described in the Methods (n = 9 WT and 9 PV δcKO mice). A trend toward a rearing difference between genotypes was detected (F (1, 16) = 3.422, P=0.0829). **C)** The possible motor coordination phenotype was further evaluated with a multi-trial accelerating rotarod test. No genotype difference was discerned (n = 9 WT and 9 PV δcKO mice; F (1, 16) = 1.703, P=0.2103). **D)** Similarly, mice performed indistinguishably on an inverted screen test (n = 9 WT and 9 PV δcKO mice, P >0.99, Mann-Whitney test). **E)** and **F)**. Cognitive screening was performed with an alternating Y-maze test. No difference in alternation rate of distance traveled was detected (n = 9 WT and 9 PV δcKO mice, unpaired two-tailed t tests, P = 0.66 and 0.70 respectively).

### Viral rescue of Gabrd expression in PV δcKO mice reduces presence of high amplitude bursts in REM

Because our recordings are performed in adult mice we cannot differentiate phenotypic changes arising from an absence of δ-mediated GABA signaling during the recording session from either a developmental alteration of network organization or a homeostatic compensation following *Gabrd* loss in PV+ cells. To test whether the high amplitude events observed during REM resulted from lack of δ containing receptors during the recordings, we utilized a viral strategy to reintroduce *Gabrd* expression in a PV+ specific manner in adult mice (Fig. 7A). We produced AAV particles packaged in the PHP.eB serotype containing an expression construct to allow for the widespread cre-dependent expression of *Gabrd* in PV+ cells following systemic administration of virus. Retro-orbital injection of either *Gabrd*-IRES-GFP expressing “Rescue” virus or GFP expressing “Control” virus into adult PV δcKO littermates served as the primary rescue arm of the experiment. We included an additional group of PVCre^+/-^ mice with WT *Gabrd* alleles to serve as controls for possible overexpression of *Gabrd* in PV+ interneurons and potential ectopic *Gabrd* expression as not all PV+ cells in the CNS normally express *Gabr*d (Belelli and Lambert, 2005). Cell type selective viral expression in the cortex was confirmed histologically in cortical tissue through colocalization of virally expressed GFP and parvalbumin in cortical interneurons (Fig. 7B). In 5 animals examined, transduction efficiency, measured as the percentage of PV-immunopositive cells that were positive for GFP, was 79.6 ± 4.6%. Selectivity, measured as the percentage of GFP-positive cells that were PV-immunopositive, was 95.7 ± 2.2%. We did observe expression of GFP in thalamic nuclei other than the PV+ reticular nucleus, but we attribute this expression to low PV expression in this population, as indicated by the Allen Brain Atlas (Yao et al., 2021).

**Figure 7.**
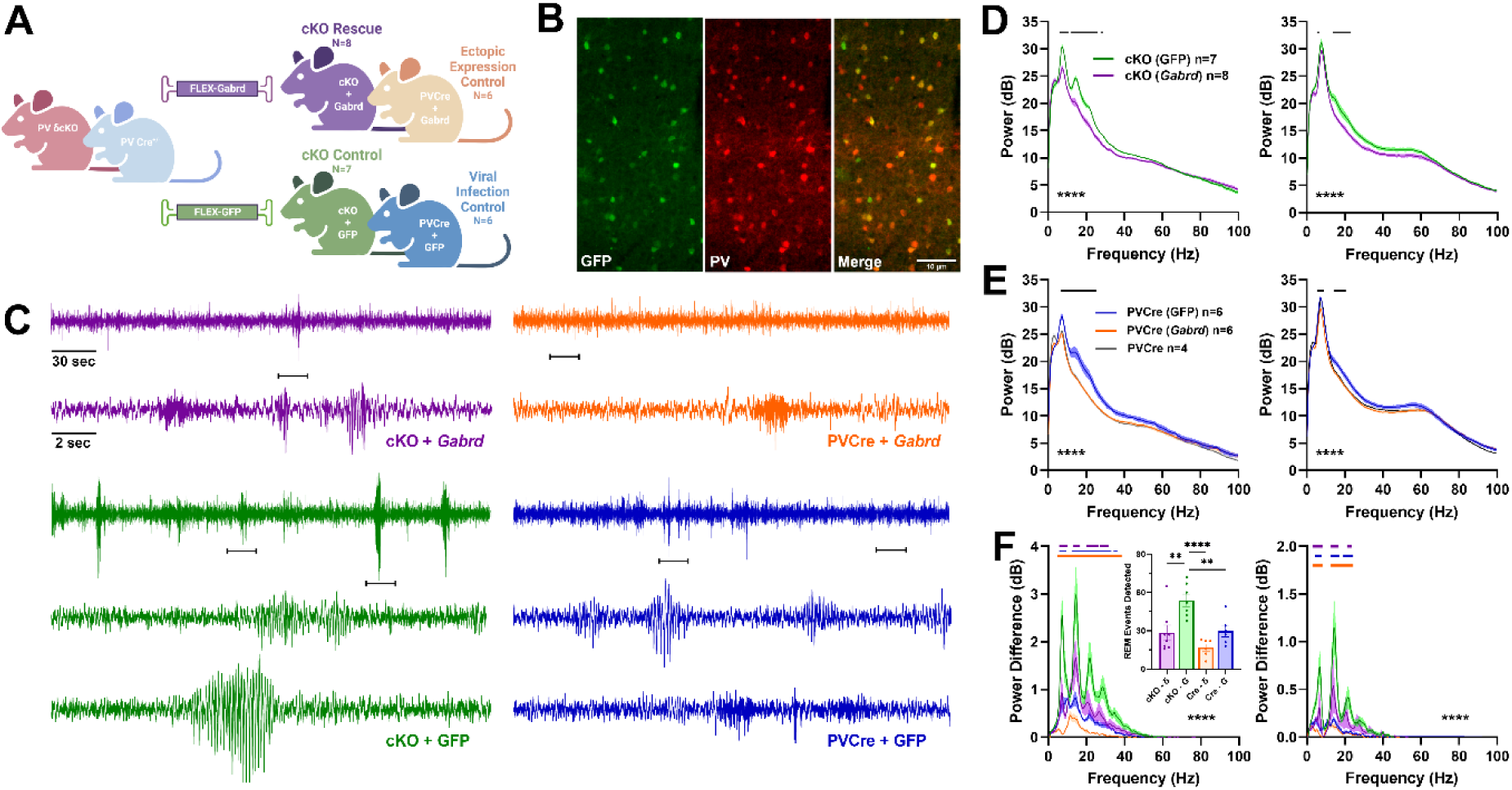
Viral delivery of Gabrd to PV δcKO mice reduces high amplitude bursts of theta-frequency oscillations during REM sleep. **A)** Experimental design for viral reintroduction of *Gabrd* into PV δcKO mice. Mice of either PV δcKO (*Gabrd*^fl/fl^ X PVCre^+/-^) or PVCre^+/-^ background were injected systemically with AAV.PhP.eB particles containing either cre-dependent *Gabrd-*IRES-GFP or GFP expression cassette. **B)** Co-expression of viral GFP with parvalbumin (PV) antibody staining in cortex indicating cell type specificity of viral expression pattern **C)** Five minutes of combined frontal REM EEG from a group of viral treated mice. Horizontal brackets on records for each mouse indicate regions shown on expanded timescale. Fewer, and lower amplitude events are observed in PVδ cKO mice that received *Gabrd* (purple) than those that were treated with GFP control (green). All records are on the same amplitude scale to demonstrate difference in the size of events during REM. **D)** REM power spectra in viral treated PV δcKO mice show significant Treatment X Frequency interaction for both frontal (left, F (255, 3328) = 8.288, p<0.0001) and parietal (right, F (255, 3328) = 1.806, p<0.0001) spectra. **E)** REM power spectra in PVCre^+/-^ control mice show significant Treatment X Frequency interaction for both frontal (left, F (510, 3328) = 2.839, p<0.0001), and parietal (right, F (510, 3328) = 1.876, p<0.0001) EEG. **F)** REM event spectra from frontal (left) and parietal (right) electrodes indicate a significant reduction in the prominence of high amplitude transient activity in PV δcKO mice treated with *Gabrd* expressing virus compared with GFP (G) control. Two Way ANOVA revealed significant Treatment X Frequency interaction for both frontal (F (765, 5888) = 5.910, p<0.0001), and parietal (F (765, 5888) = 4.056, p<0.0001) EEG. *inset*: Total number of events detected during REM, significant interaction from One-Way ANOVA (F (3, 23) = 10.03, p=0.0002). Horizontal bars in panels D and E indicate frequencies with significant group differences following Šídák’s multiple comparisons test, bars and asterisks in panel F represent frequencies that are different from the PV δcKO mice treated with GFP virus following Dunnett’s multiple comparisons.

Following a four-week incubation period after injection to allow for adequate expression of viral constructs, we again performed 12-hour light cycle EEG recordings to assess changes to the transient high amplitude events in REM following the reintroduction of *Gabrd* into PV+ cells of PV δcKO mice. In pooled REM states across the recording session, abnormal events were reduced in PV δcKO mice that received rescue virus yet persisted in their littermates that received control viral injections (Fig. 7C left records). These results support the primary hypothesis that abnormalities arise from changes to acute GABA signaling through δ-containing receptors in PV neurons rather than from a developmental consequence of early gene deletion.

We did observe some unexpected results in other limbs of the experiment. Typical REM EEG was observed in PVCre^+/-^ mice that received the rescue viral construct, suggesting no strong phenotype was associated with δ overexpression. When compared to PV^+/-^ mice that received rescue virus, those that were treated with control GFP expressing virus showed some unexpected differences in REM spectra (Fig. 7E purple), but they remained free of the transient high amplitude, long-duration events observed in PV δcKO mice (Fig. 7C right records).

In PV δcKO mice, average power spectra from REM states showed a loss of multi-peak signature in frontal EEG in the rescue condition (Fig. 7D, left). However, in both frontal and parietal EEG a broader separation between rescue and GFP conditions was present in beta and low gamma frequencies that was not previously observed between WT and PV δcKO mice. Interestingly, similar differences were present between the two viral conditions in PVCre^+/-^ mice (Fig. 7E). We interpret these results to suggest a common unanticipated effect of the GFP viral condition across both genetic backgrounds. Additionally, PV^+/-^ mice in the rescue condition showed no gross differences from naive PVCre^+/+^ mice (Fig. 7E) supporting a lack of over/ectopic expression effects by the rescue construct. Finally, event spectra, generated as described above, showed multi-peaked signature remained in PV δcKO mice injected with control virus, and the power of these peaks was reduced in all other conditions in both frontal and parietal EEG (Fig. 7F). These results support the conclusion that GFP-associated effects described above were not associated with high-amplitude REM events in PVCre^+/-^ δcKO mice.

Sleep scoring of the viral cohort showed an unexpected higher proportion of time spent in NREM in both groups receiving GFP virus, driven by increased NREM bout duration (8.16 ± 0.27 min (pooled rescue), 10.41 ± 0.51 min (pooled GFP), p=0.0005 unpaired t-test). However, no difference in REM bout count or average REM duration was detected.

Overall, the findings demonstrate the effective rescue of normal REM EEG signatures by viral reintroduction of *Gabrd* through attenuation of high amplitude transient events in PV δcKO mice, providing valuable insights into the role of *Gabrd* in PV+ neurons for regulating oscillatory activity during REM states, an effect not anticipated by previous work on the role of PV neurons in brain oscillations.

## Discussion

GABA_A_Rs containing δ subunits have been primarily studied in select principal cell types (Nusser et al., 1998; Stell and Mody, 2002; Cope et al., 2005; Brickley and Mody, 2012). PV+ interneurons also express functional δ subunit containing GABA_A_Rs, yet their role in controlling PV+ related brain activity remains unclear (Glykys et al., 2007). After confirming selective functional loss *Gabrd* in PV+ cells, we surveyed the impact on cortical EEG. PV δcKO mice exhibited similar sleep and waking behavior to WT littermates, but PV δcKO mice exhibited altered power spectra of parietal and especially frontal EEG during both NREM and REM sleep states. Increased sigma frequency oscillations in PV δcKO mice corresponded to higher amplitude and longer sleep spindles with no increase in spindle number. During REM sleep, PV δcKO mice displayed transient bursts of oscillations with a principal frequency near 7 Hz. REM spectral analysis revealed a multipeak signature in PV δcKO mice with higher power in frontal than parietal EEG. Finally, viral reintroduction of *Gabrd* expression into PV+ cells of PV δcKO mice rescued the altered REM EEG. Our results demonstrate an important acute role for slow inhibition in PV+ neurons for the maintenance of normal EEG structures of REM sleep.

Previous studies of PV δcKO mice examined hippocampal gamma oscillations *in vitro* and *in vivo* (Ferando and Mody, 2013, 2015; Barth et al., 2014). Constitutive *Gabrd* KO mice show a higher frequency peak of kainate-induced *in vitro* CA3 gamma oscillations, and both heterozygous and homozygous PV δcKO mice showed a higher peak frequency than WT animals (Ferando and Mody, 2013, 2015). An *in vivo* study of CA1 LFPs demonstrated that ovarian cycle-linked fluctuation of gamma power was abolishedin female PV δcKO mice (Barth et al., 2014). Our study provides a broader EEG survey of cortical network activity. The design included frontal EEG, which exhibited the largest sleep-associated phenotypic differences in oscillatory activities. Increased spindle amplitudes and durations in PV δcKO mice could indicate more cells responding to ascending thalamic inputs during spindles, as PV+ cortical interneuron activity increases during spindle events (Niethard et al., 2018; Brécier et al., 2022), and absence of δ mediated inhibition of these cells should increase their recruitment by excitatory inputs.

The emergence of transient high amplitude events during REM states was unexpected based previously reported subtle REM phenotypes in hippocampal LFP from PV δcKO mice (Barth et al., 2014). The events in PV δcKO mice have similar frequencies and topographic distributions to spike-wave discharges (SWDs) in models of absence epilepsy (Coenen et al., 1991; Frankel et al., 2005). Although the events were relatively suppressed during active wake, when they occurred during quiet wakefulness they predicted behavioral arrest, possibly consistent with an absence seizure. *Gabrd* mutations are associated with human epilepsies (Dibbens et al., 2004; Macdonald et al., 2012). Therefore, future study of waking behavior in these mice could be investigated as a novel model of absence epilepsy. However, the events in PV δcKO mice diverge from typical discharges seen in absence epilepsy models due to occurrence in REM. REM sleep in both rodent models and patients with absence epilepsy is typically void of SWD (Strohl et al., 2007; Ng and Pavlova, 2013). Furthermore, SWD epileptiform activity that develops in APP/PS1 mice spares REM (Jin et al., 2018).

The phenotype also shares similarity to narcolepsy. Hypocretin (HCRT)/orexin deficient narcoleptic mice exhibit remarkably similar transient events of high amplitude 7 Hz bursts of activity during REM sleep (Bastianini et al., 2012; Vassalli et al., 2013). Events in HCRT KO mice are exaggerated in frontal EEG and occur in medial prefrontal cortex but not hippocampus (Vassalli et al., 2013). If high amplitude REM events in PV δcKO mice result from similar circuit changes as REM events in HCRT KO mice, then previous PV δcKO studies utilizing hippocampal LFP would likely not detect these events (Vassalli et al., 2013; Barth et al., 2014). Interestingly, the EEG phenotype was not associated with deficits in the behavioral studies we conducted (Fig. 6).

Cortical PV+ interneurons exhibit high levels of activity during REM sleep (Niethard et al., 2018; Aime et al., 2022; Brécier et al., 2022). Although EEG analysis prevents determining whether the high amplitude events observed during REM correlate with aberrant PV+ interneuron activity *in vivo*, our rescue experiments support a role for δ mediated inhibitory tone in PV+ cells for preventing this abnormal activity. It is unclear why the loss of slow/tonic inhibition of PV+ neurons would manifest selectively in REM sleep. Changes to the ambient GABA concentrations across brain states could activate extrasynaptic δ-subunit containing GABA_A_Rs. Tonic current in hippocampus and cortex is regulated by synaptic activity (Glykys et al., 2007; Trujeque-Ramos et al., 2018); however, dynamic regulation of GABA transporters (Gaspary et al., 1998; Richerson and Wu, 2003) and direct release by astrocytes (Liu et al., 2000; Kozlov et al., 2006) are also potential mechanisms that could regulate ambient GABA. A better understanding of the dynamics of extracellular GABA across brain states will help clarify how the loss of tonic inhibition in PV+ neurons may result in phenotypes limited to specific brain states.

While *Gabrd* expression in PV+ cells has only been reported in interneurons of the cortex, hippocampus, and amygdala (Glykys et al., 2007; Ferando and Mody, 2015; Yao et al., 2021; Antonoudiou et al., 2022), our cre/lox approach ablates *Gabrd* in all PV cells across the brain. Additionally, our choice of widespread systemic delivery for viral rescue doesn’t allow for regional or PV subtype specificity. Therefore, we cannot exclude potential effects of PV δcKO in cell populations beyond those that motivated our study. Non-cortical PV neurons relevant to oscillations herein include those found in the reticular nucleus of thalamus (nRT) whose activity drives the transition of thalamocortical cell firing mode to NREM burst firing and spindle oscillations (Fernandez and Lüthi, 2020). Considerable δ-associated tonic inhibition occurs in the thalamus, yet nRT PV+ cells do not respond to delta selective agonists or express δ subunits (Pirker et al., 2000; Cope et al., 2005). Therefore, it is unlikely that ablation of *Gabrd* in nRT neurons alone drives the oscillatory phenotypes presented here.

Additionally, *Gabrd* expressing thalamic relay nuclei of the ventrobasal complex and the lateral dorsal nucleus transcribe PV RNA, yet protein expression in these regions is below the threshold of detection by immunohistochemical methods (Tanahira et al., 2009). Moreover, PV cre driver lines drive reporter genes in these regions (Tanahira et al., 2009). Although these nuclei are not classically considered PV+, the activity of the PV locus in these cells would delete *Gabrd*. We observe evidence of cre activity in these cells demonstrated by expression of viral GFP in our study, raising concern for off target cKO of *Gabrd* in a portion of thalamocortical cells. Although signaling through δ-containing receptors in thalamic relay nuclei has been implicated in electrocortical features of slow wave sleep (Vyazovskiy et al., 2005; Mesbah-Oskui et al., 2014), the thalamic nuclei affected by unintended cre activity project to cortical regions distant from the frontal EEG electrodes that show the largest phenotype (Vertes et al., 2015),.

Further, constitutive KO of *Gabrd*, which removes tonic inhibitory currents from all thalamocortical cells, does not cause abnormal high amplitude events in EEG during REM sleep (Mesbah-Oskui et al., 2014).

Our interpretation of the extent of rescue following viral reintroduction of *Gabrd* into PV δcKO mice is complicated by additional EEG findings in animals receiving GFP virus. However, despite a common increase in power of beta and low gamma ranges in GFP treated mice, high amplitude REM events remained reliably elevated and detectable only in PV δcKO and not PVCre^+/-^ mice in this viral condition. The basis for GFP-induced changes is not clear, but we note that GFP expression levels, judged by fluorescence, were much higher in control conditions than in the δ subunit rescue condition. Thus, we suspect that the unexpected effects of overexpression of GFP did not influence the rescue arm of experiments (Fig. 7). Despite these limitations, the present study demonstrates a novel role for δ subunit containing GABA_A_Rs on PV+ neurons in the regulation of normal oscillations during sleep states. Future studies to determine what cellular and circuit activities underlie the high amplitude REM events observed in PV δcKO will be a crucial step to elucidate the mechanism by which tonic inhibition of PV neurons contributes to the maintenance of normal REM oscillations in the cortex.

## Conflict of Interest

The work was funded by NIMH grants MH123748 (SM), MH122379 (CFZ, SM), MH126548 (PML), NICHD grant P50 HD103525 (Washington University Intellectual and Developmental Disability Research Center), the Taylor Family Institute for Innovative Psychiatric Research (SM, CFZ) and the Bantly Foundation (CFZ). CFZ is a member of the Scientific Advisory Board for Sage Therapeutics and holds equity in Sage Therapeutics. Sage Therapeutics had no role in the design or interpretation of the experiments herein. The remaining authors declare no competing financial interests.

## Acknowledgements

The authors thank members of the Taylor Family Institute for Innovative Psychiatric Research for discussion and input. We thank Dr. Jamie Maguire for the floxed δ mice, Dr. Jin-Moo Lee for the PV-Cre mice, and Sara Conyers for assistance with behavior data collection.

## Author contributions

Conceptual study design: PML, SVS, XL, CY & SM. Data collection and analysis: PL, SVS, XL, AB, CL, HJS, & NR. Original draft: PML. Critical revisions: PML, SVS, NR, MW, CFZ, CY, & SM.

